# Moving in pain - A preliminary study evaluating the immediate effects of experimental knee pain on locomotor biomechanics

**DOI:** 10.1101/2024.04.12.589293

**Authors:** Jesse M. Charlton, Elyott Chang, Sabrina W. Hou, Ernest Lo, Emily McClure, Cole Plater, Samantha Wong, Michael A. Hunt

**Affiliations:** School of Kinesiology, Faculty of Education, The University of British Columbia; School of Biomedical Engineering, Faculty of Medicine, The University of British Columbia; Motion Analysis and Biofeedback Laboratory, The University of British Columbia; Graduate Programs in Rehabilitation Sciences, Faculty of Medicine, The University of British Columbia; Department of Physical Therapy, Faculty of Medicine, The University of British Columbia

**Author notes:** **Corresponding Author:** Dr Jesse Charlton, PhD Postdoctoral Fellow School of Kinesiology, The University of British Columbia, Canada E.

**Keywords:** Gait, Biomechanics, Experimental Pain, Knee

## Abstract

Pain changes how we move, but it is often confounded by other factors due to disease or injury. Experimental pain offers an opportunity to isolate the independent affect of pain on movement. We used cutaneous electrical stimulation to induce experimental knee pain during locomotion to study the short-term motor adaptions to pain. While other models of experimental pain have been used in locomotion, they lack the ability to modulate pain in real-time. Twelve healthy adults completed the single data collection session where they experienced six pain intensity conditions (0.5, 1, 2, 3, 4, 5 out of 10) and two pain delivery modes (tonic and phasic). Electrodes were placed over the lateral infrapatellar fat pad and medial tibial condyle to deliver the 10 Hz pure sinusoid via a constant current electrical stimulator. Pain intensity was calibrated prior to each walking bout based on the target intensity and was recorded using an 11-point numerical rating scale. Knee joint angles and moments were recorded over the walking bouts and summarized in waveform and discrete outcomes to be compared with baseline walking. Knee joint angles changed during the swing phase of gait, with higher pain intensities resulting in greater knee flexion angles. Minimal changes in joint moments were observed but there was a consistent pattern of decreasing joint stiffness with increasing pain intensity. Habituation was limited across the 30-90 second walking bouts and the electrical current needed to deliver the target pain intensities showed a positive linear relationship. Experimental knee pain shows subtle biomechanical changes and favourable habituation patterns over short walking bouts. Further exploration of this model is needed in real-world walking conditions and over longer timeframes to quantify motor adaptations.

## Introduction

The mechanical, structural, and neurophysiological factors associated with musculoskeletal pathologies, like knee osteoarthritis, are known to alter motor control and can be observed during locomotion [1-3]. Pain is thought to be a key driver of these changes [4, 5]. However, it is difficult to parse the independent contributions of pain given that the signs and symptoms of clinical populations vary in time, intensity, and character while being interrelated with the mechanical, structural, and neurophysiological factors of the condition.

Experimental pain models isolate the experience of pain from typical disease-specific factors. While many techniques have been developed over the last half century or more [6-10], hypertonic saline injection is often used when studying knee pain. Injections into thigh muscles or the anterior knee can cause reduced internal joint moments and small changes in kinematics during walking [11-15]. Unfortunately, saline injection has important limitations, including difficulty controlling or modulated pain intensity and the use of needles that could generate unexpected responses [16-18]. A less invasive and more controllable method would provide greater flexibility.

Cutaneous electrical stimulation can provide a convenient experimental pain method that addresses the limited controllability and individualization of saline injection. Experimental pain intensity can be readily modulated based on the amplitude of electrical current [19-22], allowing for rapid changes in pain perceptions. Therein, pain can be induced based on movement mechanics in real time, such as plantar foot pressure during gait [23], ground reaction forces during standing balance [24], or wrist kinematics [20]. A limitation of this method is habituation with repeated or constant stimulation [25], but it appears this can be mitigated in the short term by using low frequency sinusoids [21, 24, 26]. Two recent studies have demonstrated that these low frequency sinusoidal stimulation protocols can induce moderate knee pain along with observable neuromuscular and mechanical adaptions [24, 27]. However, these studies were performed in static conditions (standing, seated), so it unknown if these findings extend to dynamic conditions like locomotion.

Our overarching objective was to perform an exploratory investigation of locomotor biomechanics with and without experimental pain. We aimed to 1) identify the pain intensities that create significant changes in biomechanics relative to pain-free walking; 2) evaluate changes over the course of a single walking bout; and 3) examine whether walking under a phasic stimulation pattern (linked to the gait cycle) produces different biomechanics when compared to tonic stimulation.

## Materials and methods

Drawing from Henriksen, Graven-Nielsen [13], the simple effect size for the late stance knee adduction moment was *d*=1.35 between control and experimental knee pain conditions. For a repeated measures analysis of variance with 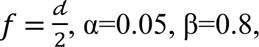 1 group, 7 measurements, and correlation among measurements of 0.5 we needed four participants. Given that we are testing on a treadmill (not over-ground) and using a different experimental pain method, we chose to recruit three times this estimate.

Twelve healthy adults were recruited between April and August 2023 from the surrounding University population to participate in this study. We included individuals who reported having no lower limb musculoskeletal injuries or surgeries in the previous 6 months, no pain in the lower limbs, or an inability to walk unaided. The study consisted of a within-subject, single-session examination of perceived pain intensity and the biomechanical changes associated with acute experimental knee pain during locomotion. The methods were approved by the Institutional Ethics Review Board (H22-03504) and the volunteers provided written informed consent prior to enrolling in the study.

Prior to beginning the experimental trials, participants completed the Pain Catastrophizing Scale questionnaire to quantify their thoughts regarding painful experiences [28]. This scale sums scores (0 = not at all, 4 = all the time) across 13 statements that represent the degree to which a person experiences various thoughts or feelings related to pain. A higher score represents more intense catastrophizing thoughts and feelings. Participants were also familiarized with verbally reporting pain intensity on a 0-10 point scale (0 = “no pain at all” and 10 = “worst pain imaginable”), which we used to monitor perceived pain intensity throughout the study.

A total of ten walking trials were collected, including a baseline trial (30 seconds in duration) with no experimental pain and nine experimental pain trials with increasing pain intensities (Figure 1). After the baseline walking trial, six phasic pain conditions were performed with increasing target pain intensities (0.5, 1, 2, 3, 4, 5 out of 10) followed by three tonic pain conditions (1, 3, 5 out of 10). Walking trials under phasic pain lasted 90 seconds, while trials under tonic pain lasted 30 seconds. All the walking conditions were completed in the participants’ personal shoes and at a self-selected speed while on a motorized, instrumented treadmill. The walking speed was established during a familiarization period lasting approximate 2-5 minutes at the start of the session. This speed was held constant throughout the subsequent experimental trials.

**Figure 1.**
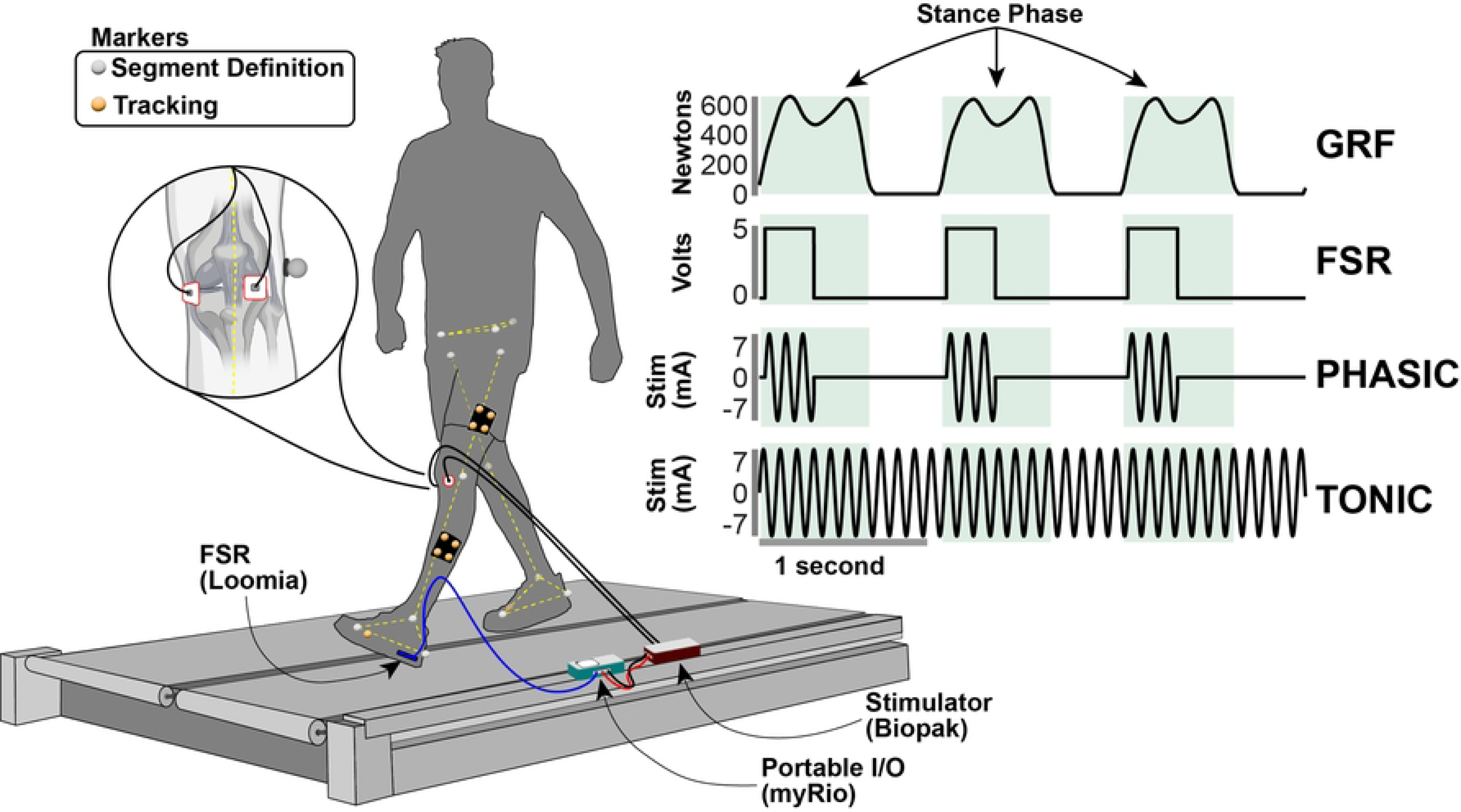
Experimental Setup. Participants walked on an instrumented treadmill while markers placed on the body were tracked using motin capture cameras. Grey markers represent segment definition markers used to construct the lower body model, while orange markers were used to track the segments with six-degree-of-freedom. A force senstive resistor (FSR) was placed inside the shoe and under the study limb heel to trigger the stimulation during the phasic trials. During the tonic trials, the sitmulation was constant. The surface electrodes were placed over the lateral infrapatellar fatpad and the medial tibial condyle (see inset of knee). The vertical ground reaction force waveform (GRF) and the green regions are shown for referencing the stance phase of the gait cycle with respect to the FSR and stimulation signals.

### Experimental Pain

Electrical stimuli were delivered via surface electrodes (3M 2560, 4 x 3.5 cm, MN, USA) placed on the skin over the lateral infrapatellar fat pad and the medial tibial condyle of the left knee (cathode on the medial location). The left knee was selected as the study limb due to equipment and space limitations. The electrodes were present on the skin for all testing conditions, even if no electrical stimulation was applied. The infrapatellar fat pad is well innervated by nociceptors [29], as are structures like the periosteum [30], making these locations ideal for an experimental pain protocol. Based on pilot testing, these electrode locations produce painful sensations across the anteromedial aspects of the knee; a pain pattern with clinical relevance to symptomatic knee osteoarthritis [31, 32] and patellofemoral pain [33]. Excitation of sensory afferents with ramping electrical current (e.g., triangle, sinusoid waveform) can preferentially recruit smaller diameter and higher threshold polymodal C-fibres while bypassing larger Aδ fibres [21, 34]. Compared to what is observed with rectangular waveforms [35], this more selective activation could contribute to lower adaptation rates by limiting potential sensory gating when larger pools of afferents are stimulated [36]. Moreover, the polymodal fibres are associated with secondary, mechanical, and dull pain sensations that are common in musculoskeletal disease and injury [37].

To control the electrical stimulation, we used a portable I/O device (myRIO 1900, National Instruments, TX, USA) and custom control software developed in Labview 2019 (National Instruments, TX, USA). A pure 10 Hz sine wave was generated at 1000 Hz and scaled to deliver the desired amplitude. The digital signal was passed through an onboard digital-to-analog converter (±10 V; 12 bits at 1000 Hz) and sent to a separate constant current isolated linear electrical stimulator (STIMSOLA, Biopac Systems Inc., CA, USA) with a 1:1 volt to current ratio. Button leads connected the stimulator to the electrodes placed on the participant.

To generate a phasic electrical stimulation based on real-time movement, we used a force sensitive resistor (2.6 x 5.1 cm; range 0.2-4.5 kg; Loomia, NY, USA) similar to previous work [23]. The sensor was taped to the plantar aspect of the left heel and connected to the I/O device’s analog input channel (±5 V; 12 bits, 1000 Hz). During the phasic pain conditions, the painful electrical stimuli were only delivered when the force sensitive resistor signal was greater than 1V. If the signal was less than 1V, no electrical stimuli were passed. This induced stimulation only during the early stance phase of gait (from heel strike to mid-stance). To compare with previous experimental pain literature (which generally involved continuous/tonic pain intensities) we collected three additional conditions of tonic pain. In these conditions, the electrical stimulation amplitude was held constant for the entirety of the 30-second walking bout regardless of the signal from the force sensitive resistor.

Prior to each walking bout we identified the electrical current needed to induce the target pain intensity (0.5, 1, 2, 3, 4, or 5 out of 10). While the participant was standing quietly, we incrementally increased the current amplitude (starting at 0.1 mA) by intervals of 0.2 mA and delivered bursts of stimulation in ∼2 second epochs. Participants were asked to indicate when the pain intensity reached the target. They then began walking on the treadmill with the same electrical current delivered and adjusted (if needed) to induce the target pain intensity while walking. Once confirmed, the walking bout began with the electrical stimulation. Pain intensity was then verbally reported at the end of each walking bout. Between every walking trial a break of 2-5 minutes was provided.

### Biomechanics Data Collection

The instrumented treadmill (Bertec Corp, OH, USA) measured ground reaction forces sampled at 2000 Hz. Simultaneously, 10 high-speed cameras (Motion Analysis Corp, CA, USA) recorded the trajectories of 35 retroreflective markers at 100 Hz. These data were synchronized in Cortex (v.8, Motion Analysis Corp, CA, USA). The lower body was defined by markers placed bilaterally on the 1^st^, 2^nd^, and 5^th^ metatarsals (exterior of the shoe), calcanei, malleoli, femoral epicondyles, the anterior superior iliac spines, and a single marker on the sacrum. For segment tracking purposes, we placed four rigid plastic plates each with four non-colinear markers on the shank and thigh. The markers on the medial malleoli, femoral condyles and 1^st^ metatarsal heads were removed after the static calibration was recorded. Biomechanics data were collected in 30-second epochs during the walking bouts, including the first and last 30 seconds of the 90-second long phasic pain trials, and the entirety of the 30-second long baseline and tonic pain trials.

### Data Processing

Marker trajectories and ground reaction forces were combined in Visual 3D (C-Motion, MD, USA) to calculate spatiotemporal, joint angle, and joint moment outcomes. We applied a lowpass Butterworth filter to the marker trajectories (order=4, cut off=6 Hz) and ground reaction force data (order=4, cut off =10 Hz). A six-degree of freedom kinematic model was used and joint angles were calculated as the distal with respect to the proximal segment. Net joint moments were calculated using an inverse dynamics approach, and were expressed as external moments, resolved in the proximal segment’s reference frame, and normalized to body mass. Positive knee angles and moments represented flexion and adduction. Joint angle and moment waveforms were time normalized to 100% of the gait cycle or stance phase, respectively.

Discrete outcomes were extracted from the biomechanical signals using custom MATLAB scripts (v. 2021b, Mathworks, MA, USA). All complete strides (heel strike to heel strike) that were captured within each 30-second data collection epoch were segmented based on force data (>20N threshold) and used in the subsequent analyses. Impulse was calculated prior to time normalizing the waveforms. Dynamic joint stiffness [38] was included as it represents the coupling between kinematic (knee joint angle) and kinetic (knee joint moment) aspects of joint motion, and has relevance in clinical populations like those with knee osteoarthritis [2, 39]. Spatiotemporal measures were included to examine the global locomotor effects in addition to the knee biomechanics outcomes. See Table 1 for specific outcomes and their definitions.

**Table 1.**
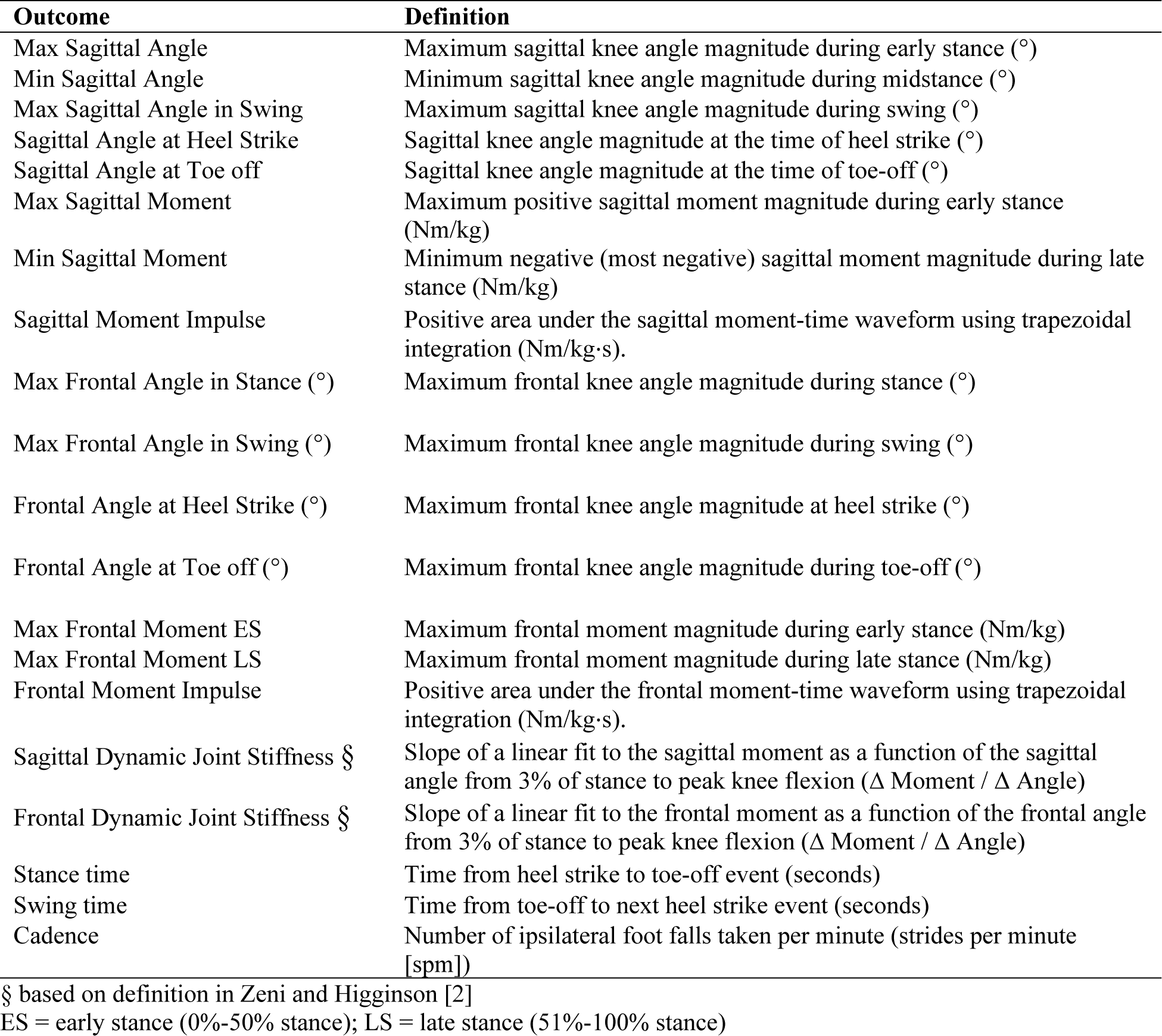
Discrete outcome definitions.

### Statistical Analysis

Discrete variables are presented as mean (standard deviation: SD) unless otherwise indicated. The following statistical tests were performed in R Stats version 4.0.3 and mainly used the *rstatix* package [40]. To address our hypotheses, we used the Wilcoxon signed-rank test to examine within-subject differences for three sets of planned comparisons. We separately tested the differences between the baseline walking bout and both the first and last 30 seconds of walking in each pain condition (pain intensities: 0.5, 1, 2, 3, 4, 5). We also compared the first 30 seconds of walking during the phasic pain conditions with the 30-second tonic pain conditions (only pain intensities of 1, 3, and 5). The effect size *r* was calculated for each signed-rank test as 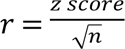 where *n* is the number of pairs which was 12. As the goal of the study was to explore the potential differences in biomechanics that could arise from experimental pain, we did not adjust the alpha level for multiplicity, which remained at 0.05 for all comparisons.

Our overarching secondary goal was to characterize the experience of pain during walking. First, pain intensity at the start and end of each walking trial was evaluated as a function of electrical current (mA) across all the pain intensity conditions. A linear model was fit to the data to quantify the increase in electrical current necessary to elicit a 1.0 unit change in pain intensity. Second, changes in perceived pain magnitude were investigated by comparing the difference in pain intensities reported at the end (following completion of the 90-second walking trial) versus the start of each walking bout via a two-sided Student’s t-test (non-equal variance).

## Results

The twelve participants in our study were on average 27 (4) years old, 1.76 (0.12) m in body height, had a body mass of 73.1 (17.7) kg, were all right limb dominant, and five indicated their sex at birth as female. The median total score for the pain catastrophizing scale was 3 (interquartile: 2, 16), which suggests generally low catastrophizing thoughts and feelings amongst our study sample. One participant did not complete the tonic pain conditions and two participants’ data were omitted from comparisons involving joint moments due to a calibration error between the force plates and motion capture frame of reference.

### Effects of Experimental Pain (Phasic and Tonic)

In general, the phasic and tonic pain conditions resulted in several small changes in knee joint biomechanics (Figure 2) when compared to the baseline walking condition, predominantly in the sagittal plane (Table 2). During swing, participants increased their knee flexion angle by approximately 1-1.5° which was significantly different in the 2, 3, and 4 out of 10 pain intensity conditions (*r*>0.61, *p*<0.034) but not for the 5 out of 10 condition (*r*=0.52, *p*=0.077). However, knee flexion angle in swing during the final 30 seconds of data collection remained elevated in the 3 out of 10 condition (*r*=0.61, *p*=0.034) and the 5 out of 10 condition became significant (*r*=0.86, *p*=0.001). All three tonic pain conditions elicited higher knee flexion angles in swing (*r*>0.69, *p*<0.027). The only other kinematic difference was knee flexion angle at toe-off, which was greater for the 3 out of 10 condition during the first 30 seconds, the 1 and 2 out of 10 conditions during the last 30 seconds, and the 1 and 3 out of 10 tonic pain conditions (*r*>0.61, *p*<0.034). As can be seen in Figure 2, the frontal plane angle during swing appeared to show a consistent shift toward higher adduction angles with higher pain intensity. While the variance in the sample was too large to detect significant differences (*p*>0.52), this may require further investigation. Otherwise, we did not observe any frontal plane angle differences that were significant (Table 3).

**Figure 2.**
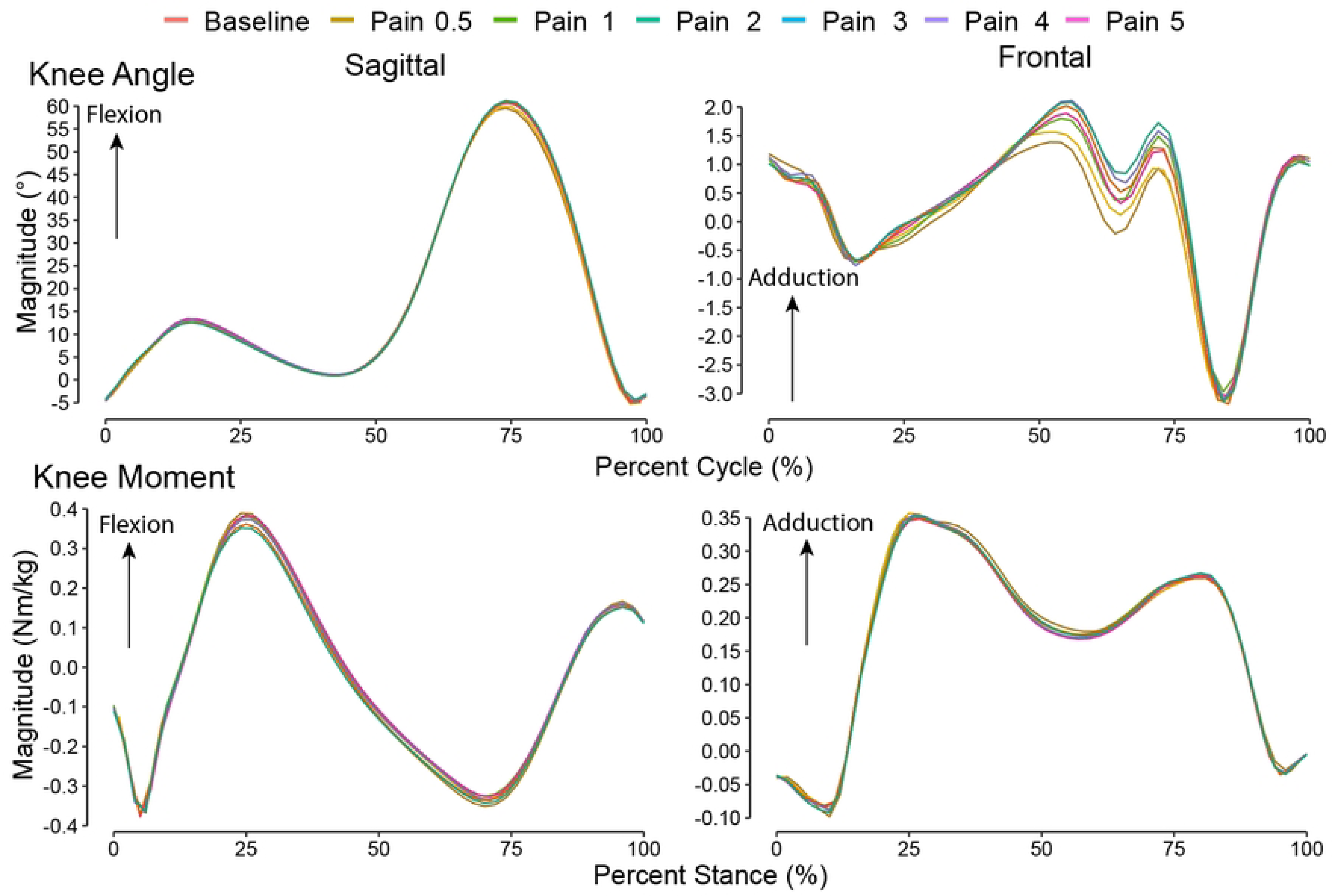
Knee joint waveforms. Sagittal and frontal knee joint biomechanics as ensemble average waveforms from all participants in the experimental pain conditions (phasic and tonic combined for visualization).

**Table 2.** Discrete sagittal plane biomechanics outcomes. Mean (SD) for each condition (Baseline, Phasic Pain, Tonic Pain) at each pain level (0.5, 1, 2, 3, 4, 5) extracted from the start (0 to 30 seconds [0:30s]) and the end (60 to 90 seconds [60:90s]) of the walking bout. Joint moment outcomes were calculated on n=10.

|  | Baseline | Phasic, 0.5 |  | Phasic, 1 |  | Phasic, 2 |  | Phasic, 3 |  | Phasic, 4 |  | Phasic, 5 |  | Tonic, 1 | Tonic, 3 | Tonic, 5 |
| --- | --- | --- | --- | --- | --- | --- | --- | --- | --- | --- | --- | --- | --- | --- | --- | --- |
| Outcome | 0:30s | 0:30s | 60:90s | 0:30s | 60:90s | 0:30s | 60:90s | 0:30s | 60:90s | 0:30s | 60:90s | 0:30s | 60:90s | 0:30s | 0:30s | 0:30s |
| Max Sagittal Angle in Stance (°) | 13.33<br>(5.25) | 13.08<br>(5.57) | 13.10<br>(5.60) | 13.09<br>(5.33) | 13.31<br>(5.59) | 13.63<br>(5.74) | 13.50<br>(5.88) | 13.30<br>(6.42) | 13.13<br>(6.12) | 12.77<br>(6.58) | 12.11<br>(5.20) | 12.32<br>(6.30) | 12.36<br>(6.15) | 14.18<br>(5.38) | 14.13<br>(5.31) | 14.07<br>(5.38) |
| Min Sagittal Angle in Stance (°) | -7.12<br>(4.38) | -7.88<br>(5.77) | -7.89<br>(4.92) | -7.58<br>(5.41) | -7.39<br>(5.27) | -6.89<br>(5.63) | -7.35<br>(5.39) | -7.40<br>(5.82) | -8.11<br>(5.70) | -6.99<br>(5.12) | -7.21<br>(5.15) | -6.86<br>(4.90) | -6.98<br>(5.22) | -6.22<br>(4.91) | -6.28<br>(4.21) | -5.79<br>(4.58) |
| Max Sagittal Angle in Swing (°) | 59.74<br>(5.99) | 59.88<br>(5.89) | 60.17<br>(5.76) | 60.06<br>(5.07) | 60.82<br>(5.50) | 60.84<br>(5.46) | 60.98<br>(6.34) | 60.88<br>(5.46) | 61.06<br>(5.54) | 61.25<br>(5.61) | 59.67<br>(6.17) | 60.87<br>(6.47) | 61.71<br>(5.73) | 62.59<br>(4.81) | 62.35<br>(4.45) | 62.50<br>(4.56) |
| Sagittal Angle at Heel Strike (°) | -3.61<br>(4.41) | -3.74<br>(5.18) | -3.79<br>(5.24) | -3.55<br>(5.11) | -3.31<br>(5.96) | -3.46<br>(6.00) | -3.79<br>(5.32) | -3.31<br>(6.11) | -3.73<br>(5.27) | -3.33<br>(5.85) | -3.51<br>(4.98) | -3.11<br>(5.44) | -3.49<br>(5.55) | -2.80<br>(5.40) | -2.64<br>(5.29) | -2.27<br>(5.61) |
| Sagittal Angle at Toe off (°) | 38.09<br>(5.14) | 38.93<br>(4.95) | 38.47<br>(5.82) | 38.44<br>(5.99) | 40.07<br>(6.86) | 39.08<br>(5.69) | 39.65<br>(5.91) | 39.66<br>(5.83) | 38.99<br>(6.76) | 39.00<br>(6.27) | 38.44<br>(5.41) | 39.44<br>(6.51) | 39.57<br>(6.59) | 41.28<br>(4.61) | 40.84<br>(4.14) | 40.28<br>(4.93) |
| Max Sagittal Moment (Nm/kg) | 0.42<br>(0.23) | 0.42<br>(0.23) | 0.42<br>(0.22) | 0.42<br>(0.22) | 0.42<br>(0.24) | 0.40<br>(0.20) | 0.41<br>(0.22) | 0.41<br>(0.24) | 0.39<br>(0.21) | 0.38<br>(0.22) | 0.38<br>(0.20) | 0.39<br>(0.22) | 0.38<br>(0.23) | 0.42<br>(0.26) | 0.42<br>(0.25) | 0.41<br>(0.25) |
| Min Sagittal Moment (Nm/kg) | -0.44<br>(0.10) | -0.44<br>(0.10) | -0.45<br>(0.10) | -0.44<br>(0.10) | -0.45<br>(0.10) | -0.45<br>(0.10) | -0.46<br>(0.10) | -0.47<br>(0.10) | -0.46<br>(0.09) | -0.45<br>(0.10) | -0.45<br>(0.13) | -0.46<br>(0.10) | -0.45<br>(0.10) | -0.45<br>(0.09) | -0.46<br>(0.10) | -0.45<br>(0.12) |
| Sagittal Moment Impulse (Nm/kg-s) | 0.07<br>(0.04) | 0.07<br>(0.05) | 0.07<br>(0.04) | 0.07<br>(0.05) | 0.07<br>(0.05) | 0.06<br>(0.04) | 0.07<br>(0.05) | 0.07<br>(0.05) | 0.06<br>(0.04) | 0.06<br>(0.05) | 0.06<br>(0.04) | 0.06<br>(0.04) | 0.06<br>(0.04) | 0.07<br>(0.06) | 0.07<br>(0.05) | 0.07<br>(0.06) |
| Sagittal Dynamic Joint Stiffness ($\Delta$ Moment / $\Delta$ Angle) | 0.039<br>(0.015) | 0.036<br>(0.013) | 0.037<br>(0.017) | 0.035<br>(0.018) | 0.032<br>(0.021) | 0.030<br>(0.011) | 0.026<br>(0.013) | 0.034<br>(0.020) | 0.026<br>(0.010) | 0.028<br>(0.012) | 0.031<br>(0.018) | 0.022<br>(0.013) | 0.024<br>(0.009) | 0.031<br>(0.018) | 0.031<br>(0.016) | 0.031<br>(0.007) |

*ES, early stance (0%-50% stance); LS, late stance (51%-100% stance). Positive values are flexion, adduction, and greater dynamic stiffness*

**Table 3.** Discrete frontal plane biomechanics outcomes. Mean (SD) for each condition (Baseline, Phasic Pain, Tonic Pain) at each pain level (0.5, 1, 2, 3, 4, 5) extracted from the start (0 to 30 seconds [0:30s]) and the end (60 to 90 seconds [60:90s]) of the walking bout. Joint moment outcomes were calculated on n=10.

|  | Baseline | Phasic, 0.5 |  | Phasic, 1 |  | Phasic, 2 |  | Phasic, 3 |  | Phasic, 4 |  | Phasic, 5 |  | Tonic, 1 | Tonic, 3 | Tonic, 5 |
| --- | --- | --- | --- | --- | --- | --- | --- | --- | --- | --- | --- | --- | --- | --- | --- | --- |
| Outcome | 0:30s | 0:30s | 60:90s | 0:30s | 60:90s | 0:30s | 60:90s | 0:30s | 60:90s | 0:30s | 60:90s | 0:30s | 60:90s | 0:30s | 0:30s | 0:30s |
| Max Frontal Angle in Stance (°) | 3.26<br>(2.08) | 3.57<br>(1.94) | 3.27<br>(2.24) | 3.42<br>(2.25) | 3.57<br>(2.30) | 3.63<br>(2.05) | 3.45<br>(2.17) | 3.82<br>(2.02) | 3.70<br>(2.22) | 3.67<br>(2.25) | 3.63<br>(2.21) | 3.77<br>(2.12) | 3.80<br>(2.16) | 3.77<br>(2.76) | 3.73<br>(2.90) | 3.97<br>(2.62) |
| Max Frontal Angle in Swing (°) | 4.67<br>(2.62) | 4.88<br>(2.76) | 4.73<br>(3.02) | 4.73<br>(3.00) | 4.84<br>(3.17) | 5.04<br>(2.91) | 4.96<br>(2.87) | 5.28<br>(3.01) | 5.22<br>(3.20) | 5.05<br>(3.10) | 4.96<br>(2.74) | 5.32<br>(3.09) | 5.38<br>(2.96) | 5.54<br>(3.95) | 5.65<br>(4.32) | 5.87<br>(3.97) |
| Frontal Angle at Heel Strike (°) | 1.13<br>(2.91) | 0.98<br>(2.91) | 0.97<br>(2.79) | 0.88<br>(2.86) | 1.07<br>(2.74) | 1.16<br>(2.77) | 0.97<br>(2.88) | 1.04<br>(2.65) | 1.07<br>(2.71) | 1.09<br>(2.68) | 0.90<br>(2.79) | 0.76<br>(2.78) | 0.78<br>(2.95) | 1.13<br>(2.64) | 1.02<br>(2.64) | 1.15<br>(2.73) |
| Frontal Angle at Toe off (°) | -0.16<br>(3.85) | 0.29<br>(3.56) | 0.20<br>(3.70) | 0.33<br>(3.89) | 0.60<br>(4.00) | 0.68<br>(3.69) | 0.30<br>(3.90) | 0.84<br>(3.90) | 0.83<br>(3.97) | 0.75<br>(3.86) | 0.69<br>(4.13) | 0.86<br>(4.13) | 0.82<br>(4.12) | 1.07<br>(4.58) | 1.14<br>(4.70) | 1.44<br>(4.46) |
| Max Frontal Moment ES (Nm/kg) | 0.39<br>(0.09) | 0.39<br>(0.09) | 0.38<br>(0.09) | 0.38<br>(0.10) | 0.39<br>(0.10) | 0.38<br>(0.10) | 0.38<br>(0.10) | 0.39<br>(0.09) | 0.38<br>(0.09) | 0.38<br>(0.09) | 0.38<br>(0.09) | 0.39<br>(0.09) | 0.39<br>(0.09) | 0.37<br>(0.10) | 0.37<br>(0.11) | 0.36<br>(0.11) |
| Max Frontal Moment LS (Nm/kg) | 0.28<br>(0.10) | 0.29<br>(0.10) | 0.29<br>(0.10) | 0.28<br>(0.10) | 0.30<br>(0.11) | 0.28<br>(0.10) | 0.29<br>(0.09) | 0.29<br>(0.10) | 0.29<br>(0.10) | 0.28<br>(0.10) | 0.27<br>(0.09) | 0.29<br>(0.10) | 0.30<br>(0.10) | 0.28<br>(0.09) | 0.28<br>(0.10) | 0.28<br>(0.10) |
| Frontal Moment Impulse (Nm/kg·s) | 0.13<br>(0.04) | 0.13<br>(0.04) | 0.13<br>(0.04) | 0.13<br>(0.04) | 0.13<br>(0.04) | 0.12<br>(0.04) | 0.13<br>(0.03) | 0.13<br>(0.04) | 0.13<br>(0.04) | 0.13<br>(0.04) | 0.12<br>(0.03) | 0.13<br>(0.04) | 0.13<br>(0.04) | 0.12<br>(0.03) | 0.12<br>(0.04) | 0.12<br>(0.04) |
| Frontal Dynamic Joint Stiffness (Δ Moment / Δ Angle) | -0.004<br>(0.011) | -0.002<br>(0.007) | -0.002<br>(0.006) | -0.004<br>(0.006) | -0.001<br>(0.004) | -0.006<br>(0.015) | -0.002<br>(0.006) | -0.002<br>(0.004) | -0.005<br>(0.010) | -0.007<br>(0.019) | -0.001<br>(0.006) | -0.007<br>(0.011) | -0.002<br>(0.007) | -0.004<br>(0.010) | -0.002<br>(0.007) | 0.000<br>(0.004) |

ES, early stance (0%-50% stance; LS, late stance (51%-100% stance). Positive values are flexion, adduction, and greater dynamic stiffness

We detected few differences in kinetic outcomes across the conditions. During late stance, we saw larger (more negative) minimum sagittal knee moment (corresponding to an external knee extension moment) in the 3 out of 10 condition (*r*=0.63, *p*=0.049) during the first 30 seconds of walking, while the maximum frontal knee moment magnitude (corresponding to an external knee adduction moment) was larger during the last 30 seconds for the 1 and 5 out of 10 conditions (*r*>0.67, *p*<0.037).

Despite the few differences in joint angles and moments, we found significant reductions in sagittal dynamic joint stiffness as a result of phasic pain compared to baseline. Over the initial 30 seconds of walking with 2, 4, and 5 out of 10 pain, participants walked with lower stiffness (*r*>0.63, *p*<0.049) which remained significantly lower in the final 30 seconds for the 2 and 5 out of 10 conditions (*r*>0.82, *p*<0.006). The 3 out of 10 condition also elicited lower stiffness in the final 30 seconds (*r*=0.79, *p*=0.01). Meanwhile, only the 5 out of 10 tonic condition elicited significantly lower stiffness (*r*=0.69, *p*=0.039). The frontal dynamic joint stiffness was not significantly different across pain intensities with the smallest p-value at 0.193 for first 30 seconds at 5 out of 10 pain relative to baseline.

The only spatiotemporal changes that were detected as a result of experimental pain were in stance time. During the first 30 seconds of the 3 and 5 out of 10 phasic conditions, we observed an increase in stance time of approximately 10 milliseconds (*r*>0.77, *p*<0.005). However, during the final 30 seconds of walking, every phasic pain condition except for 0.5 out of 10 showed an increase in stance time (*r*>0.72, *p*<0.009) between 9 and 14 milliseconds (Table 4). For tonic pain compared to baseline, the 1 and 3 out of 10 conditions had longer stance times of approximately 9-10 milliseconds (*r*>0.69, *p*<0.027).

**Table 4.** Discrete spatiotemporal biomechanics outcomes. Mean (SD) for each condition (Baseline, Phasic Pain, Tonic Pain) at each pain level (0.5, 1, 2, 3, 4, 5) extracted from the start (0 to 30 seconds [0:30s]) and the end (60 to 90 seconds [60:90s]) of the walking bout.

|  | Baseline | Phasic, 0.5 |  | Phasic, 1 |  | Phasic, 2 |  | Phasic, 3 |  | Phasic, 4 |  | Phasic, 5 |  | Tonic, 1 | Tonic, 3 | Tonic, 5 |
| --- | --- | --- | --- | --- | --- | --- | --- | --- | --- | --- | --- | --- | --- | --- | --- | --- |
| Outcome | 0:30s | 0:30s | 60:90s | 0:30s | 60:90s | 0:30s | 60:90s | 0:30s | 60:90s | 0:30s | 60:90s | 0:30s | 60:90s | 0:30s | 0:30s | 0:30s |
| Stance Time (s) | 0.690<br>(0.046) | 0.693<br>(0.051) | 0.694<br>(0.051) | 0.694<br>(0.049) | 0.704<br>(0.048) | 0.692<br>(0.054) | 0.699<br>(0.052) | 0.700<br>(0.049) | 0.700<br>(0.049) | 0.693<br>(0.052) | 0.704<br>(0.047) | 0.700<br>(0.051) | 0.704<br>(0.048) | 0.694<br>(0.049) | 0.693<br>(0.043) | 0.691<br>(0.045) |
| Swing Time (s) | 0.414<br>(0.024) | 0.407<br>(0.023) | 0.412<br>(0.024) | 0.409<br>(0.029) | 0.406<br>(0.028) | 0.408<br>(0.027) | 0.407<br>(0.029) | 0.406<br>(0.031) | 0.412<br>(0.032) | 0.412<br>(0.033) | 0.410<br>(0.032) | 0.408<br>(0.030) | 0.409<br>(0.034) | 0.404<br>(0.031) | 0.409<br>(0.033) | 0.413<br>(0.034) |
| Cadence (strides/min) | 54.5<br>(3.2) | 54.8<br>(3.5) | 54.5<br>(3.5) | 54.7<br>(3.6) | 54.3<br>(3.3) | 54.8<br>(3.8) | 54.5<br>(3.9) | 54.6<br>(3.8) | 54.1<br>(3.7) | 54.6<br>(4.0) | 54.2<br>(3.8) | 54.5<br>(3.8) | 54.1<br>(3.6) | 54.9<br>(3.6) | 54.7<br>(3.5) | 54.6<br>(3.7) |

### Phasic versus Tonic Pain

We compared the phasic and tonic pain conditions to examine possible differences in immediate motor response based on stimulation delivery method. The primary finding was that the tonic pain conditions generally resulted in larger joint angle magnitudes. Maximum knee flexion magnitude was higher in the 5 out of 10 tonic condition compared to the phasic condition (*r*=0.62, *p*=0.042); while the 1 out of 10 tonic condition resulted in higher knee flexion at toe-off and during swing (*r*>0.67, *p*<0.024). Lastly, when compared to the phasic condition, the 5 out of 10 tonic condition produced higher sagittal dynamic joint stiffness (*r*=0.73, *p*=0.020). No frontal plane outcomes were different between phasic and tonic pain conditions. Swing time was the only significant spatiotemporal difference, with the 5 out of 10 condition resulting in a 7 millisecond shorter swing time in the tonic condition compared to the phasic (*r*=0.64, *p*=0.032).

### Pain Intensity

The relationship between the electrical current and the perception of pain intensity showed a positive relationship (Figure 3). The relationship could be approximated by a linear fit which indicated that a 1-unit increase in pain required a 0.64-0.67 mA increase in our study sample.

**Figure 3.**
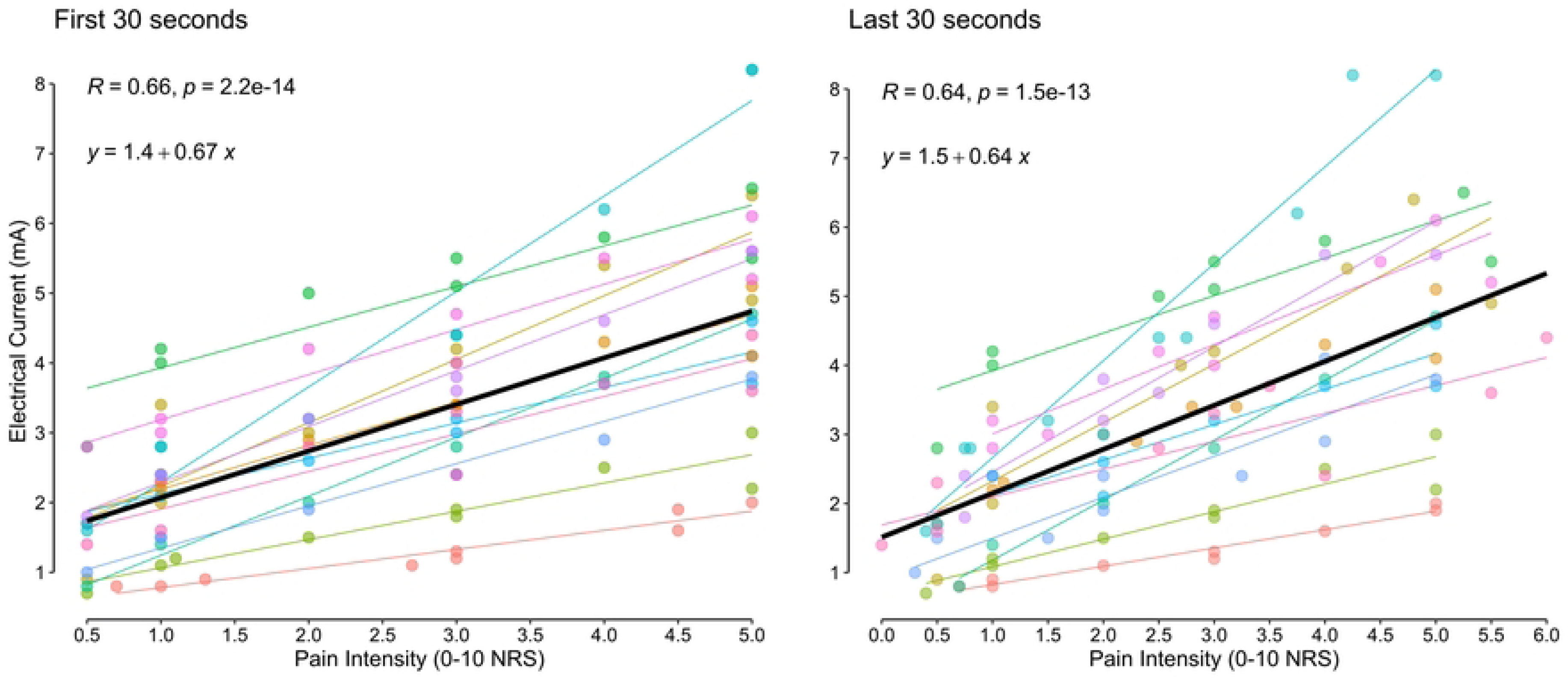
Electrical current as a function of pain. A linear model (black line = all participants, coloured lines = individual participants) was fit to the reported pain intensity and electrical current data for each walking condition. On average a 1-point increase in pain required an increase of 0.67 mA and 0.64 mA, based on the first and last 30 seconds of walking, respectively.

On average, pain intensity was stable from the start to the end of each phasic walking bout. However, individual responses varied, as can be seen by the data points included in Figure 4, with some showing evidence of sensitizing (increased magnitude) and others habituating (decreased magnitude) over the walking bout. The tonic pain condition showed similar patterns except for the 5 out of 10 condition which appeared to show a significant sensitization across our study sample (*p*=0.016).

**Figure 4.**
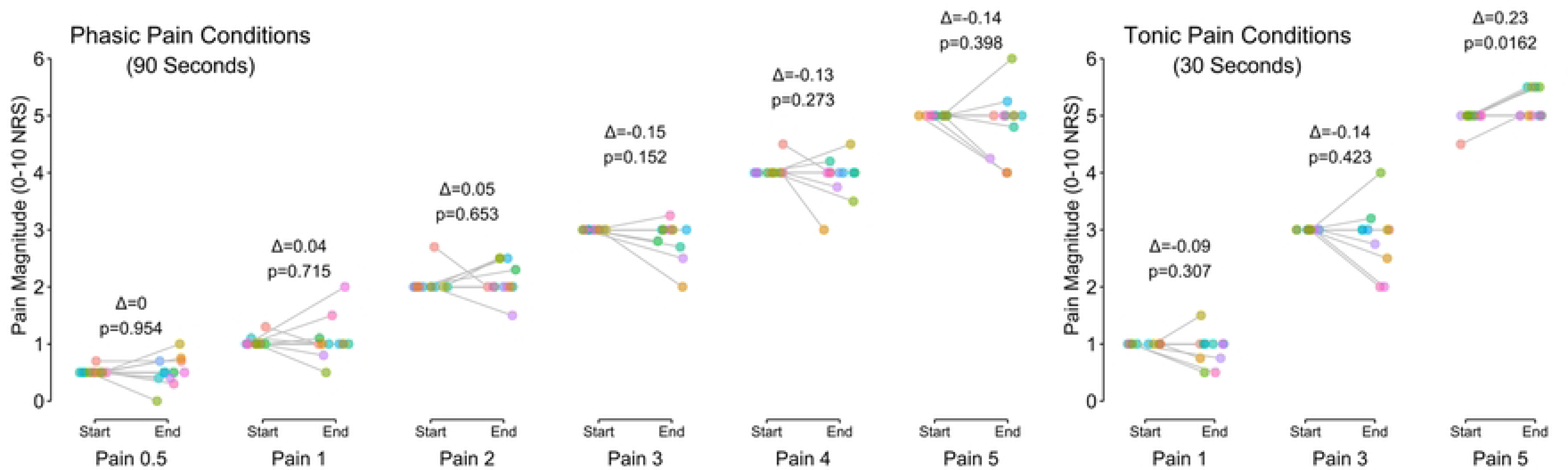
Pain reports. At the start and end of each experimental condition, participants reported their perception of pain on an 11-point NRS. Grey dots are individual participants connected by lines between the start and end of their trial. Annotated results show the mean change (Δ) and p-value of the Student’s t-test.

## Discussion

Experimental pain models offer the opportunity to investigate the relationships between pain and motor control. Based on the advantageous properties of cutaneous electrical stimulation, we generated painful stimuli at incremental intensities that were linked to the gait cycle (phasic) or constant over time (tonic). The biomechanical responses to different pain intensities were subtle in our study, including higher knee flexion angles and lower dynamic sagittal plane joint stiffness, depending on the intensity of pain. Meanwhile few differences were observed between the phasic and tonic pain delivery methods for a given pain intensity. Our use of a treadmill may have limited the changes in biomechanics due to pain given that speed was fixed. The relationship between pain intensity and electrical current could be approximated by a linear function and we did not observe habituation or sensitization, on average, in all but one condition. Together, this study brings new contributions including identifying changes in dynamic joint stiffness, assessing the feasibility of using experimental pain modulation during gait, and establishing the relationship between electrical stimulation amplitude and pain intensity during locomotion.

Previous experimental knee pain studies have reported different knee kinematics during locomotion, based on the location that the pain was induced. With hypertonic saline injection in the quadriceps, sagittal knee angles were largely unaffected [11], while anterior knee injections resulted in 1.5° reductions in maximum sagittal knee angles during stance compared to no pain [13]. We did not observe a significant change in maximum sagittal knee angles during stance, though a small, but noticeable, reduction can be seen in the group mean data. Nevertheless, the kinematic changes during stance were small in our study and similarly so in previous literature, which suggests knee kinematics are not sensitive to the presence of experimental pain.

The more consistent kinematic change that we observed during the experimental conditions was an increase in swing phase knee flexion angles. These results loosely support, and are of a similar magnitude to, results from treadmill running while participants received short-duration electrical square-wave pulses to induce pain at varying time-points of the gait cycle [22]. In a later study, stimulation was presented at 4 different times over the gait cycle and knee flexion increases during swing were also observed, although unlike our study the swing phase increases appear to be a result of late stance stimulation [41]. It is possible that this occurs due to a reduction in quadriceps muscle activation from experimental anterior knee pain [42]. A flexion-reflex response as an explanation is less likely, given the similar findings between phasic and tonic stimulation in our study and the kinematic responses being later than the expected delays of 125-300ms [41, 43]. Further research is needed to ascertain the mechanisms of these changes.

When comparing the kinetic results from our study to previous literature that used different experimental pain modalities, two differences stand out. First, we observed few changes in general, and the outcomes that were significantly different were small in magnitude. In past work, hypertonic saline injections in the anterior knee generated significant reductions relative to the control for both peak sagittal (∼0.03 Nm/kg) and frontal (∼0.02-0.08 Nm/kg) plane joint moments [13]. Our mean differences in the higher pain intensities for maximum sagittal joint moments (0.04 Nm/kg) were of a similar order, though we saw smaller changes in peak frontal moments (0.01 Nm/kg). The reductions we did see may be the result of changes in trunk position, like a forward lean, given the limited changes in knee angles during stance. This action would reduce the sagittal moment arm about the knee joint center of rotation and lower the flexion moment. We did not measure trunk kinematics and so this explanation remains as speculation.

Dynamic joint stiffness describes the relationship between changes in moments about the knee with respect to the concomitant changes in joint angle. It is common to see increases in dynamic joint stiffness in clinical populations like those with knee osteoarthritis [2, 44, 45]. Given the structural knee joint changes that occur in these populations, it is possible that the increase in stiffness is driven largely by the musculoskeletal features, neuromuscular adaptions associated with the condition, or the presence or absence of pain in general, and less so by the intensity of any pain experienced. Our results suggest that experiencing pain generates larger reductions in peak sagittal knee moments relative to the changes in maximum knee joint angles during early stance, which would generate smaller dynamic joint stiffness magnitudes. As our sample has normal knee function, this result could be a product of responding to the pain conditions with upper body/postural changes that affect the ground reaction force but not the knee kinematics directly (as described earlier).

Despite different total exposure to the electrical stimulation between phasic and tonic conditions, we saw few biomechanical differences. With limited data to compare with, two possible explanations could be considered. First, the salience of both phasic and tonic stimulation was similar enough such that the motor responses were generally comparable. If we assume the perceived pain intensity as analogous to salience, our methods would have predicted this outcome given that we calibrated each condition individually. Second, the overall exposure time and movement constraints imposed by the treadmill may have resulted in minimal motor adaptation in general. It is possible that more readily measurable changes in biomechanics could be observed during over-ground walking when spatiotemporal features of gait can be more readily modulated.

We were able to induce target pain intensities by modulating the electrical current prior to beginning the walking bouts. Following previous reports [7, 20, 21] we found that increasing electrical stimulation intensity generated increased pain perception. This relationship between electrical current and pain perception was linear, like findings in previous studies that involved upper limb pain stimulation [20, 21, 46]. Although there were clear individual differences in the current required to induce a given pain intensity, the positive relationship was consistent for pain reports at the beginning of the walking bout (Figure 3, left panel) and similar but more variable after the walking bout (Figure 3, right panel). This likely reflects the different habituation/sensitization trajectories we observed for each participant, resulting in a different pain rating at the end of each walking bout (Figure 4).

While there were individual differences in pain reports over the walking bouts on average there was minimal habituation or sensitization. Gallina, Abboud [24] used the same protocol as we did and found stable pain intensity ratings over 60 seconds during tonic stimulation while sitting quietly. However, square waves and higher frequency sinusoids showed more consistent habituation patterns (reduction in pain intensity) of approximately 1 point on the 11-point NRS [24]. Our data support these past results, while extending the evidence for limited pain perception changes over 90 seconds and while walking.

We found that there were differences between the required electrical stimulation amplitude necessary to induce target pain intensities between the stimulation conditions. Comparing between phasic and tonic pain, we found that 6 (55%), 7 (64%), and 10 (91%) participants required lower electrical current during the tonic conditions. This amounted to an average -0.06 (0.68) mA, -0.45 (0.85) mA and -1.09 (1.27) mA reduction in current to elicit the target pain intensity, respectively. The need for more current in the phasic condition could be related to limited temporal summation due to the shorter period of exposure to a single stimulation burst. With phasic pain being triggered by the force sensitive resistor underfoot, the stimuli lasted approximately 300-400ms (this was not recorded), but could take 2-3x that time to reach a temporal summation plateau [47].

Several limitations need to be considered to further contextualize our findings. We only applied the experimental pain condition for a short time and longer exposure may have resulted in different biomechanical responses. Participants walked on a treadmill and were unable to modify their gait speed in response to the experimental pain conditions. It is plausible that participants would have chosen different gait speeds if able, which would likely have required differences in kinematics and kinetic outcomes. Continued development of this experimental pain method should include conditions where volitional changes in spatiotemporal gait characteristics are permitted. The experimental pain conditions were presented in a non-randomized order of increasing intensity. This was to avoid any potential carryover of high intensity conditions to the lower intensity conditions or possible ‘startle’ like effects when going from baseline to higher pain intensities.

The impact of pain on motor control and movement biomechanics continues to be an important area of research where experimental pain models have significant utility. One objective is to establish how and why we move differently when in pain, in the hopes of providing more targeted and individualized treatment for those living with painful conditions. Experimental pain models provide a means of studying this in a controlled fashion. We applied electrical stimulation to the knee joint during locomotion to study changes in biomechanics and pain perceptions. Small changes in knee biomechanics were observed and predominantly occurred in the sagittal plane, including higher knee flexion angles and lower dynamic joint stiffness. Pain perceptions were generally stable across the exposures. The key advancement of this model is the ability to have pain and movement coupled in real-time. This will support investigations of how movement and pain relationships are adapted to during discrete or continuous tasks.

## References

1. Petterson SC, Barrance P, Buchanan T, Binder-Macleod S, Snyder-Mackler L. Mechanisms underlying quadriceps weakness in knee osteoarthritis. Medicine and Science in Sports and Exercise. 2008;40(3):422–7.

2. Zeni JA, Higginson JS. Dynamic knee joint stiffness in subjects with a progressive increase in severity of knee osteoarthritis. Clinical Biomechanics. 2009;24(4):366–71.

3. Hutchison L, Grayson J, Hiller C, D’Souza N, Kobayashi S, Simic M. Relationship between knee biomechanics and pain in people with knee osteoarthritis: A systematic review and meta-analysis. Arthritis Care & Research. 2023;75(6):1351–61.

4. Hodges PW, Coppieters MW, MacDonald D, Cholewicki J. New insight into motor adaptation to pain revealed by a combination of modelling and empirical approaches. European Journal of Pain. 2013;17(8):1138–46.

5. Hodges PW, Tucker K. Moving differently in pain: A new theory to explain the adaptation to pain. PAIN. 2011;152(3).

6. Wolf S, Hardy JD. Studies on pain. Observations on pain due to local cooling and on factors involved in the "cold pressor" effect. J Clin Invest. 1941;20(5):521–33.

7. Rainville P, Feine JS, Bushnell MC, Duncan GH. A psychophysical comparison of sensory and affective responses to four modalities of experimental pain. Somatosens Mot Res. 1992;9(4):265–77.

8. Steinbrocker O, Isenberg SA, Silver M, Neustadt D, Kuhn P, Schittone M. Observations on pain produced by injection of hypertonic saline into muscles and other supportive tissues. J Clin Invest. 1953;32(10):1045–51.

9. Woolf CJ. Transcutaneous electrical nerve stimulation and the reaction to experimental pain in human subjects. PAIN. 1979;7(2):115–27.

10. Bank PJ, Peper CE, Marinus J, Beek PJ, van Hilten JJ. Motor consequences of experimentally induced limb pain: a systematic review. European Journal of Pain. 2013;17(2):145–57.

11. Henriksen M, Alkjaer T, Lund H, Simonsen EB, Graven-Nielsen T, Danneskiold-Samsoe B, et al. Experimental quadriceps muscle pain impairs knee joint control during walking. Journal of Applied Physiology. 2007;103:132–9.

12. Henriksen M, Rosager S, Aaboe J, Bliddal H. Adaptations in the gait pattern with experimental hamstring pain. Journal of Electromyography and Kinesiology. 2011;21(5):746–53.

13. Henriksen M, Graven-Nielsen T, Aaboe J, Andriacchi TP, Bliddal H. Gait changes in patients with knee osteoarthritis are replicated by experimental knee pain. Arthritis Care & Research. 2010;62(4):501–9.

14. Simonsen MB, Yurtsever A, Næsborg-Andersen K, Leutscher PDC, Hørslev-Petersen K, Andersen MS, et al. Tibialis posterior muscle pain effects on hip, knee and ankle gait mechanics. Human Movement Science. 2019;66:98–108.

15. Seeley MK, Park J, King D, Hopkins JT. A novel experimental knee-pain model affects perceived pain and movement biomechanics. Journal of Athletic Training. 2013;48(3):337–45.

16. Bennell K, Hodges P, Mellor R, Bexander C, Souvlis T. The nature of anterior knee pain following injection of hypertonic saline into the infrapatellar fat pad. Journal of Orthopaedic Research. 2004;22(1):116–21.

17. Graven-Nielsen T, Arendt-Nielsen L, Svensson P, Jensen TS. Experimental muscle pain: A quantitative study of local and referred pain in humans following injection of hypertonic saline. Journal of Musculoskeletal Pain. 1997;5(1):49–69.

18. Deacon B, Abramowitz J. Fear of needles and vasovagal reactions among phlebotomy patients. Journal of Anxiety Disorders. 2006;20(7):946–60.

19. Laursen RJ, Graven-Nielsen T, Jensen TS, Arendt-Nielsen L. Quantification of local and referred pain in humans induced by intramuscular electrical stimulation. European Journal of Pain. 1997;1(2):105–13.

20. Bergin M, Tucker K, Vicenzino B, Hodges PW. “Taking action” to reduce pain—Has interpretation of the motor adaptation to pain been too simplistic? PLOS ONE. 2021;16(12):e0260715.

21. Jonas R, Namer B, Stockinger L, Chisholm K, Schnakenberg M, Landmann G, et al. Tuning in C-nociceptors to reveal mechanisms in chronic neuropathic pain. Annals of neurology. 2018;83(5):945–57.

22. Duysens J, Tax AAM, Trippel M, Dietz V. Phase-dependent reversal of reflexly induced movements during human gait. Experimental Brain Research. 1992;90(2):404–14.

23. Bertrand-Charette M, Jeffrey-Gauthier R, Roy J-S, Bouyer LJ. Gait adaptation to a phase-specific nociceptive electrical stimulation applied at the ankle: A model to study musculoskeletal-like pain. Frontiers in human neuroscience. 2021;15.

24. Gallina A, Abboud J, Blouin J-S. A task-relevant experimental pain model to target motor adaptation. Journal of Physiology. 2021;599:2401–17.

25. Eitner L, Özgül ÖS, Enax-Krumova EK, Vollert J, Maier C, Höffken O. Conditioned pain modulation using painful cutaneous electrical stimulation or simply habituation? European Journal of Pain. 2018;22(7):1281–90.

26. Grill WM, Mortimer JT. Stimulus waveforms for selective neural stimulation. IEEE Engineering in Medicine and Biology Magazine. 1995;14(4):375–85.

27. Cabral HV, Devecchi V, Oxendale C, Jenkinson N, Falla D, Gallina A. Effect of movement-evoked and tonic experimental pain on muscle force production. Scandinavian Journal of Medicine & Science in Sports. 2024;34(1):e14509.

28. Sullivan MJL, Bishop SR, Pivik J. The Pain Catastrophizing Scale: Development and validation. Psychological Assessment. 1995;7(4):524–32.

29. Bohnsack M, Hurschler C, Demirtas T, Rühmann O, Stukenborg-Colsman C, Wirth CJ. Infrapatellar fat pad pressure and volume changes of the anterior compartment during knee motion: possible clinical consequences to the anterior knee pain syndrome. Knee Surg Sports Traumatol Arthrosc. 2005;13(2):135–41.

30. Steverink JG, Oostinga D, van Tol FR, van Rijen MHP, Mackaaij C, Verlinde-Schellekens SAMW, et al. Sensory innervation of human bone: An immunohistochemical study to further understand bone pain. The Journal of Pain. 2021;22(11):1385–95.

31. Felson DT. The sources of pain in knee osteoarthritis. Current Opinion in Rheumatology. 2005;17(5):624–8.

32. Creamer P, Lethbridge-Cejku M, Hochberg M. Where does it hurt? Pain localization in osteoarthritis of the knee. Osteoarthritis and Cartilage. 1998;6(5):318–23.

33. Gerbino PGI, Griffin ED, d’Hemecourt PA, Kim T, Kocher MS, Zurakowski D, et al. Patellofemoral pain syndrome: evaluation of location and intensity of pain. The Clinical Journal of Pain. 2006;22(2):154–9.

34. Accornero N, Bini G, Lenzi GL, Manfredi M. Selective activation of peripheral nerve fibre groups of different diameter by triangular shaped stimulus pulses. Journal of Physiology. 1977;273(3):539–60.

35. Milne RJ, Kay NE, Irwin RJ. Habituation to repeated painful and non-painful cutaneous stimuli: a quantitative psychophysical study. Experimental Brain Research. 1991;87(2):438–44.

36. Melzack R, Wall PD. Pain mechanisms: A new theory. 5th ed 1965. 971-9 p.

37. Nikolenko VN, Shelomentseva EM, Tsvetkova MM, Abdeeva EI, Giller DB, Babayeva JV, et al. Nociceptors: Their role in Body’s defenses, tissue specific variations and anatomical updates. J Pain Res. 2022;15:867–77.

38. Davis RB, DeLuca PA. Gait characterization via dynamic joint stiffness. Gait & Posture. 1996;4(3):224–31.

39. Chang AH, Chmiel JS, Almagor O, Guermazi A, Prasad PV, Moisio KC, et al. Association of baseline knee sagittal dynamic joint stiffness during gait and 2-year patellofemoral cartilage damage worsening in knee osteoarthritis. Osteoarthritis and Cartilage. 2017;25(2):242–8.

40. Kassambara A. rstatix: Pipe-friendly framework for basic statistical tests. 0.7.2 ed2023.

41. Spaich EG, Arendt-Nielsen L, Andersen OK. Modulation of lower limb withdrawal reflexes during gait: A topographical study. Journal of Neurophysiology. 2004;91(1):258–66.

42. Park J, Hopkins JT. Induced anterior knee pain immediately reduces involuntary and voluntary quadriceps activation. Clinical Journal of Sport Medicine. 2013;23(1):19–24.

43. Sandrini G, Serrao M, Rossi P, Romaniello A, Cruccu G, Willer JC. The lower limb flexion reflex in humans. Progress in neurobiology. 2005;77(6):353–95.

44. Gustafson JA, Anderton W, Sowa GA, Piva SR, Farrokhi S. Dynamic knee joint stiffness and contralateral knee joint loading during prolonged walking in patients with unilateral knee osteoarthritis. Gait & Posture. 2019;68:44–9.

45. McGinnis K, Snyder-Mackler L, Flowers P, Zeni J. Dynamic joint stiffness and co-contraction in subjects after total knee arthroplasty. Clinical Biomechanics. 2013;28(2):205–10.

46. Willer JC. Comparative study of perceived pain and nociceptive flexion reflex in man. PAIN. 1977;3(1):69–80.

47. Arendt-Nielsen L, Brennum J, Sindrup S, Bak P. Electrophysiological and psychophysical quantification of temporal summation in the human nociceptive system. European Journal of Applied Physiology and Occupational Physiology. 1994;68(3):266–73.

